# Foveal vision predictively sensitizes to defining features of eye movement targets

**DOI:** 10.1101/2022.01.11.475331

**Authors:** Lisa M. Kroell, Martin Rolfs

## Abstract

Despite the fovea’s singular importance for active human vision, the impact of large eye movements on foveal processing remains elusive. Building on findings from passive fixation tasks, we hypothesized that during the preparation of rapid eye movements (saccades), foveal processing anticipates soon-to-be fixated visual features. Using a dynamic large-field noise paradigm, we indeed demonstrate that sensitivity for defining features of a saccade target is enhanced in the pre-saccadic center of gaze. Enhancement manifested in higher Hit Rates for foveal probes with target-congruent orientation, and a sensitization to incidental, target-like orientation information in foveally presented noise. Enhancement was spatially confined to the center of gaze and its immediate vicinity. We suggest a crucial contribution of foveal processing to trans-saccadic visual continuity which has previously been overlooked: Foveal processing of saccade targets commences before the movement is executed and thereby enables a seamless transition once the center of gaze reaches the target.

## Introduction

Foveal processing is of singular importance for primate vision. Within a small, central region of the retina known as the foveal pit, retinal layers spread aside and allow light to impinge directly onto a densely packed population of cone photoreceptors^[1]^. The resulting signals carry highly resolved visual information, the prioritization of which is reflected in the organization of upstream neural areas: even though the fovea covers merely 0.01% of the retina, more than 8% of primary visual cortex is devoted to the processing of foveal input^[2–4]^. To utilize the resolution of those signals, primates routinely and rapidly move their eyes to bring relevant information into foveal vision.

Quite surprisingly, foveal processing appears understudied on both a neurophysiological and behavioral level^[5]^. In the moving observer in particular, decades of research have characterized pre-saccadic sensitivity modulations at the target of eye movements^[6–8]^. To date, little to nothing is known about the concurrent development of visual sensitivity in the pre-saccadic center of gaze. We hypothesized that the fovea is not merely a passive receiver of input for high-acuity vision. Instead, it appears uniquely suited to predict incoming information and, in consequence, to assume a crucial role in the establishment of visual continuity across saccades.

Evidence from neuroimaging^[9,10]^, brain stimulation^[11]^ and psychophysics^[10,12,13]^ suggests that during fixation, the fovea contributes to the processing of stimuli presented in the periphery: using functional magnetic resonance imaging, Williams et al. (2008) and Fan et al. (2016) demonstrate that relevant peripheral objects are represented in foveal retinotopic cortex, possibly via feedback connections from temporal areas^[14,15]^. Foveal cortex thus seems to be recruited for tasks requiring high perceptual scrutiny – even when the respective stimulus appears far outside any foveal neuron’s receptive field. Indeed, disrupting foveal processing through transcranial magnetic stimulation^[11]^ or the presentation of a foveal distractor^[10,13]^ impairs peripheral discrimination performance. Foveal feedback, so far characterized during fixation, gains a predictive nature when applied to active vision^[9,11]^: already before an eye movement, critical features of the peripheral target should be available for foveal processing. Once gaze has shifted, the target is foveated. In the moving observer, foveal feedback therefore anticipates incoming information and may facilitate post-saccadic target processing or gaze correction when the eyes land erroneously off-target ^[16–18]^. Most notably, this mechanism would not require predictive spatial updating to support transsaccadic continuity: irrespective of the fovea’s future location, feedback to the current center of gaze would suffice to predict the target’s features on a retinotopic scale.

To investigate this potential mechanism, we characterized the immediate perceptual signature of foveal feedback during eye movement preparation. We hypothesized that fed-back information about the saccade target combines with foveal input and thereby facilitates the detection of target-congruent, foveal feature information. On a perceptual level, this should correspond to a predictive enhancement of saccade target features in the pre-saccadic center of gaze.

The lack of knowledge on foveal processing is mostly due to methodological constraints: in neurophysiological investigations, foveal receptive fields are challenging to estimate since even in paralyzed monkeys, gaze is never fully stable. Only recent developments combining a free viewing approach with an offline reconstruction of visual input allow for these estimations^[19]^. Psychophysical investigations of foveal processing require a measure sensitive enough to reveal subtle changes in high-acuity vision on the level of behavioral responses. At the same time, the stimulus probing performance in the center of gaze must be inconspicuous enough not to interfere with saccade programming^[20,21]^. In light of these considerations, we smoothly embedded our stimuli in a dynamic stream of full-screen, 1/f noise images^[22]^ (see ***Methods***). Observers maintained fixation in the screen center while the noise images flickered at a temporal frequency of 20 Hz (**Figure 1A**). At some point, the saccade target appeared 10 degrees of visual angle (dva) to the left or right of fixation, cueing the eye movement. The target was created by filtering the background noise at the respective location to an orientation of −45° or +45° (**Supplementary Figure S1**). On 50% of the trials, a probe stimulus, that is, an additional orientation-filtered noise patch, appeared in the screen center during saccade preparation and remained visible for 50 ms. Crucially, the probe was oriented either −45° or +45° and therefore either congruently or incongruently to the orientation of the target. The conspicuity of the foveal probe could be adjusted flexibly and in small increments by varying its transparency against the background noise. After executing the saccade, observers reported if they had perceived a stimulus in the screen center (present versus absent). After a ‘present’ response, they additionally reported the perceived orientation (left versus right for −45° and +45°, respectively). We investigated whether the orientation of the saccade target influenced the foveal detection judgment.

**Figure 1.**
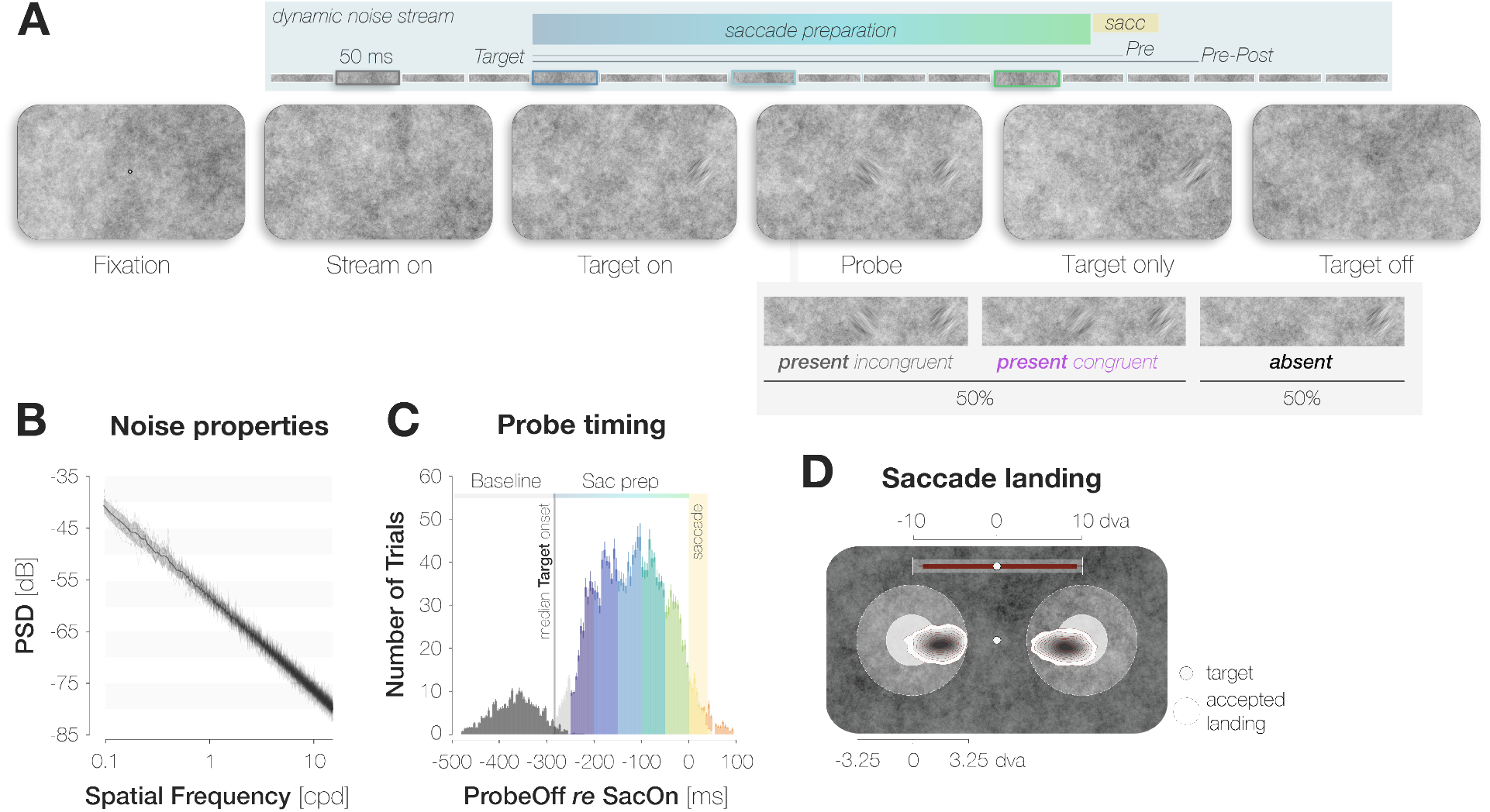
A dynamic noise paradigm probing foveal sensitivity to saccade target features. **A.** Example trial procedure in Experiment 1: The saccade target and foveal probe were embedded in full-screen noise images flickering at a frequency of 20 Hz (image duration of 50 ms). After 200 ms, the saccade target (an orientation-filtered patch; filtered to either –45° or +45°; 3 dva in diameter) appeared 10 dva to the left or the right of the screen center, cueing the eye movement. On 50% of trials, a probe (a second orientation-filtered patch; filtered to either –45° or +45°) appeared in the screen center either 150 ms before target onset (top panel; highlighted element with grey outline), or at an early (dark blue outline), medium (light blue outline) or late (green outline) stage of saccade preparation. The foveal probe was presented for 50 ms and could be oriented either congruently or incongruently to the target. On *Pre* trials, the saccade target disappeared before saccade landing. On *Pre-Post* trials, it remained visible for a brief duration after landing. Observers reported if they had perceived the probe in the screen center or not (present vs absent). After a ‘present’ response, they reported the probe’s perceived orientation (left for −45° vs right for +45°). **B.** Noise properties: Power spectral density (PSD) of the foveal region (3 dva diameter) of all noise images presented in a randomly chosen experimental session. **C.** Probe timing: histogram of time intervals between probe offset and saccade onset. Bar heights and error bars indicate the mean and standard error of the mean (SEM) across observers, respectively. On baseline trials, the probe appeared before target onset (dark grey bars). On all remaining trials, the probe appeared after target onset and therefore during saccade preparation (Sac prep). We assigned saccade preparation trials to five distinct time bins (from dark blue to light green). Trials in which the probe disappeared more than 250 ms before saccade onset (light grey), during the saccade (yellow) or after saccade offset (orange) were excluded. The yellow background rectangle illustrates the median saccade duration. **D.** Observers were able to select the peripheral stimulus as the saccade target: Bivariate Gaussian Kernel densities of saccade landing coordinates for left- and rightwards saccades. Filled circles indicate the saccade target (*rad* = 1.5 dva). Transparent circles indicate the accepted landing area (*rad* = 3.25 dva). The distance between the screen center and the targets was reduced for illustration purposes. Red bars on top indicate median saccade amplitudes based on both the horizontal and vertical component of the saccade. Neither saccade latencies nor saccade endpoints influenced congruency effects (see ‘Methods’).

We made three main observations (**Figure 2B-D**). First, observers’ responses suggest a foveal sensitization to the target’s orientation. While Hit Rates (HRs) for target-congruent and target-incongruent probes were indistinguishable long before the saccade, congruent HRs started to exceed incongruent ones 175 ms before saccade onset. In our design, the foveal region was never void of signal but contained incidental orientation information in the background noise even on probe absent trials. On those trials, we expected observers to become sensitive to target-congruent orientations in the foveal noise and, in consequence, to report the target orientation more often than the non-target orientation when generating a False Alarm (FA). Indeed, congruent False Alarm rates (FARs) exceeded incongruent ones. Second, just like the increase in HRs, the increase in FARs signifies *enhancement*. Target-congruent FAs were by no means unsystematic but relied on an incidental, high energy of the reported orientation in the foveal noise. Third, enhancement is *foveal* rather than global. Enhancement was most pronounced in the center of gaze and exhibited an asymmetric profile, extending further towards than away from the target. Based on these findings, we suggest that their predictive potential in active visual settings may constitute the key functionality of foveal feedback connections that, so far, have exclusively been characterized during passive fixation.

**Figure 2.**
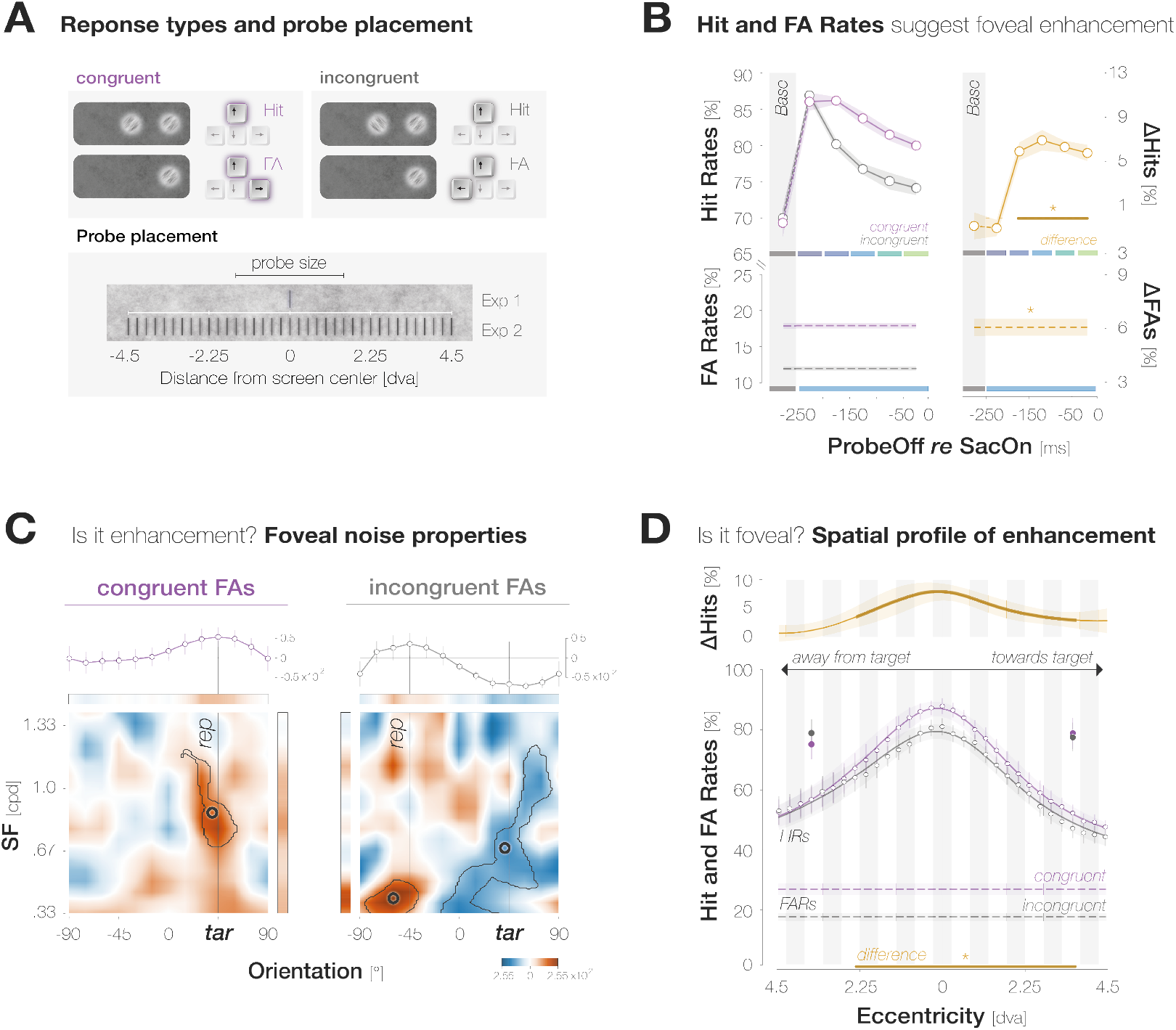
Evidence for predictive foveal enhancement of saccade target features. **A.** *Top*: Response types. (In)congruent Hits are probepresent trials in which the probe was oriented (in)congruently to the target and observers reported ‘present’ (up arrow key). (In)congruent FAs are probe-absent trials in which observers reported the presence (up arrow key) of a probe with target-(in)congruent orientation (left, right arrow keys). *Bottom*: Probe placement. In Experiment 1, we chose a single probe location in the screen center. In Experiment 2, we defined 37 probe locations spaced evenly on a horizontal axis of 9 dva length around the screen center to measure the spatial profile of enhancement. **B.** Hit and FA rates in Experiment 1 (y-axis) for different pre-saccadic time bins (x-axis; dark blue to light green rectangles), separated into congruent (purple) and incongruent (gray) responses. The difference between congruencies is plotted in brown. Asterisks indicate *p*-values <= .05. **C.** Foveal noise properties on congruent and incongruent FA trials. We described the energy of 260 combinations of orientations (x-axis) and spatial frequency (SF; y-axis) in the foveal noise region (see ***Methods***). Noise images corresponding to trials with leftward target orientation were flipped, such that +45° corresponds to the target orientation (‘tar’). The reported orientation (‘rep’) was either +45° (congruent FAs) or −45° (incongruent FAs). Orange regions indicate positive energy values in the standardized energy map (*z-score* > 0) whereas blue regions indicate negative values (*z-score* < 0). Marginal means along the orientation axis (curves and horizontal colored bars) are averages of energy values across all SFs. Marginal means along the SF axis (vertical colored bars) are averages of all SFs within the horizontal boundaries of the cluster around the reported orientation. Open circles indicate the center of mass of identified clusters. **D.** Hit and FA rates in Experiment 2 (y-axis) for locations horizontally offset from the fovea (x-axis). Note that FAs cannot be spatially resolved. Data points indicate mean response rates across observers. The plotted curves are average Gaussian function fits to individualobserver means after aligning them to the mean recorded fixation position during saccade preparation (zero on the x-axis). Negative and positive x-axis values indicate probe locations away from and towards the saccade target, respectively. Thick brown lines highlight the significant portion of the spatial difference curve (congruent HRs – incongruent HRs; top panel). Additional data points at ±3 dva eccentricity were obtained after raising peripheral performance to a foveal level by adaptively increasing probe transparency (last session of Experiment 2). All error bands and bars denote ±1 SEM.

## Results

### Hit and False Alarm rates suggest foveal enhancement of the target’s orientation

We defined congruent and incongruent HRs as the proportion of probe-present trials in which observers reported perceiving the probe, and the probe was oriented congruently and incongruently to the target, respectively (**Figure 2A**). Based on the time interval between the offset of the probe and the onset of the saccade, we assigned each Hit trial to one of five pre-saccadic time bins of 50 ms duration each (**Figure 1C**). We obtained a baseline sensitivity estimate outside the saccade preparation period by presenting the probe 150 ms *before* the saccade target on an additional subset of trials.

A two-way repeated measures ANOVA on HRs revealed significant main effects of probe offset (*F*(5,30) = 8.1, *p* < .001) and congruency (*F*(1,6) = 9.3,*p* = .023), as well as an interaction, *F*(5,30) = 3.3, *p* = .017. Irrespective of congruency, foveal HRs decreased continuously in the course of saccade preparation, when attention is known to shift to the target^[23]^ (**Figure 2B**). We observed a HR of 86.5±11.1% in the earliest pre-saccadic bin 250–200 ms before saccade onset and a significantly lower HR of 77.1±6.8% 50–0 ms before the saccade (bootstrapped *p* < .001; see ***Methods***). Congruent and incongruent HRs did not differ in the baseline (HR_cong-incong_ = - 0.7±8.5%; *p* = .589) and earliest pre-saccadic bin (HR_cong-incong_ = - 0.7±5.1%; *p* = .669). Afterwards, a stable congruency effect emerged: HRs for congruent probes significantly exceeded HRs for incongruent probes throughout the remaining bins (HR_cong-incong_ = 6.0±5.7%, 7.0±6.5%, 6.4±3.8%, 5.9±5.2%, for the time bins [–200 −150[, [–150 −100[, [–100 −50[, [–50 0[ ms, respectively; all *p*s < .003).

We defined congruent and incongruent FARs as the proportion of probe-absent trials in which observers reported perceiving the probe followed by a target-congruent and target-incongruent orientation report, respectively (**Figure 2A**). When generating a FA, observers reported the target’s orientation more often than the non-target orientation, manifesting in higher congruent than incongruent FARs (FAR_cong-incong_ = 5.9±3.3%; *p* < .001; **Figure 2B**). Note that FAs are not time-resolved since observers may have perceived the probe at any time in the course of a probe-absent trial.

In sum, the orientation of the saccade target influenced observers’ perceptual performance in their pre-saccadic center of gaze. While this suggests a predictive foveal enhancement of saccade target features, we substantiated two aspects of this assumption with further analyses and an additional experiment. First, though consistent with our hypotheses, a comparable increase in congruent HRs and FARs does not yield an advantage for congruent probes in classical sensitivity measures such as d-prime^[24]^. To ensure that FAs with target-congruent orientation report reflect an *enhancement* of orientation information in the foveal noise region, we investigated if those responses are FAs in the classical sense, that is, whether they reflect biased response behavior. Alternatively, they may constitute a systematic reaction to incidental orientation information in the foveal noise region and therefore provide further support for a sensitization to target-congruent feature information.

### Analysis of foveal noise properties supports enhancement of foveal sensitivity

We separated all FA trials by whether observers had reported the target orientation (congruent FA) or the non-target orientation (incongruent FA). On each trial, we identified all noise images that had been displayed during the potential probe presentation period, i.e., from the onset of the dynamic noise stream to saccade onset. We described the properties of the foveal noise window in each image by determining the energy of 260 combinations of orientation and spatial frequency (ori*SF) at the potential probe location (**Figure 4, *Methods***). We subsequently identified lateralized clusters of exceedingly high or low energy in the resulting energy maps using a combination of *t*-tests and bootstrapping procedures. Note that we flipped the energy maps of trials in which the target was oriented to the left such that, in all subsequent analyses and plots, +45° corresponds to the target-congruent orientation while −45° corresponds to the incongruent orientation. Details on image filtering, normalization and statistical procedures are provided in the ***Methods***.

Energy maps underlying congruent and incongruent FAs showed a clear lateralization, with regions of high energy clustering around the perceived orientation (**Figure 2C**). Noise images associated with target-congruent FAs were characterized by a high energy around the target’s orientation. The identified cluster exhibited a center of mass (CM) close to +45° (*ori_CM_* = 39.38°, *SF_CM_* = 0.86 cycles per degree (cpd); *tSum* = 6625.7; *p* < .001). Interestingly, this cluster included SFs from .67 cpd to 1.19 cpd – a range that covers the pre-saccadic development of visual resolution at the target of a 10 dva saccade^[24]^. This suggests that information about the target was indeed available for foveal processing, at a resolution influenced by the target’s pre-saccadic eccentricity. Fed-back information may have interacted with foveal noise content, allowing those ori*SF combinations that are congruent with the fed-back signal – but not others – to drive response behavior. Indeed, separating noise images that had appeared before (**Figure 3A**, ‘Baseline’) or after target onset (‘Sac prep’) revealed that the cluster encompassing higher SFs manifested exclusively during saccade preparation (*ori_CM_* = 40.31°, *SF_CM_* = 0.57 cpd; *tSum* = 22,099.0; *p* < .001).

**Figure 3.**
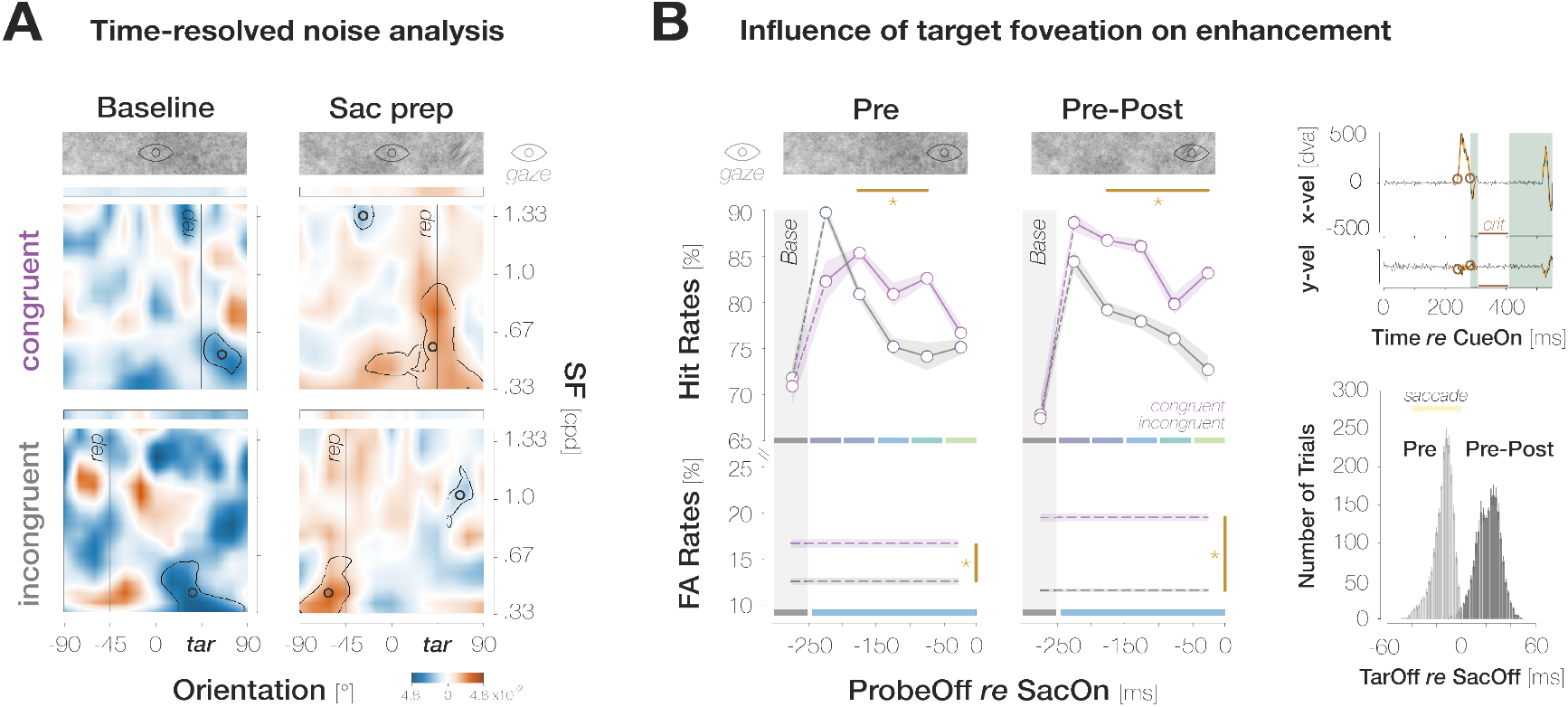
Additional findings in Experiment 1. **A.** Foveal noise properties underlying congruent (top row) and incongruent (bottom row) FAs, separately for noise images that had appeared during the baseline time bin (left column) and during saccade preparation (right column; ‘Sac prep’). All conventions follow those of Figure 2C. **B.** Main panel: Foveal Hit Rates (top row) and FA Rates (bottom row) separately for trials in which the saccade target had disappeared during saccadic flight (left column; ‘Pre’) and after the saccade (right column; ‘Pre-Post’). Conventions as in Figure 2B. *Side panel, top*: Illustration of saccade inclusion criteria based on horizontal and vertical eye velocities (x vel; y vel) recorded in an example trial. Orange outlines highlight offline-detected saccadic events. To ensure that foveal input remained stable on the retina after landing, we excluded trials in which a second saccade occurred in a critical time window (‘crit’) between 25 and 100 ms after response saccade offset. We employed a conservative estimate of saccade offset and excluded post-saccadic oscillations from the saccadic profile. We therefore introduced a short time window of 25 ms after response saccade offset during which saccadic activity did not lead to trial exclusions. *Side panel, bottom*: Histogram of time intervals between target offset and saccade offset. On *Pre* trials, the target disappeared at a median of 14.1 ms before saccade landing. On *Pre-Post* trials, the target remained visible for 22.3 ms after saccade landing.

Noise images associated with target-incongruent FAs exhibited a high energy around the perceived, non-target orientation. The corresponding cluster was slightly repelled from the target orientation (*ori_CM_* = 60.0°) and confined to very low SFs from .33 to .51 cpd (*SF_CM_* = 0.41 cpd; *tSum* = 5587.7; *p* = <.001). These SFs likely generated a salient percept when flashing on screen, motivating observers to report the non-target orientation in a purely stimulus-driven fashion. Unlike congruent FAs, incongruent FAs were not associated with a higher-SF cluster around the perceived orientation. Instead, the underlying noise images exhibited an absence of evidence for the target orientation that manifested in a cluster of low energy centered on 41.25° (*SF_CM_* = .68 cpd; *tSum* = - 20,050; *p* =.002). Whereas the low-SF portion of this cluster relied on images visible in the baseline time bin, the portion encompassing target-like orientations in a higher SF spectrum manifested exclusively during saccade preparation (**Figure 3A**). Combined, the high-energy cluster repelled from the target orientation, and the low-energy cluster covering a wide SF range around the target orientation suggest that throughout the trial, noise content as target-dissimilar as possible was required for observers to perceive a competing orientation in the foveal noise. Finally, we compared the amount of evidence required to perceive a certain orientation by summarizing the absolute filter responses within the identified clusters for each response type. Since incongruent FAs required both, perceptual evidence for the non-target orientation and an absence of evidence for the target orientation, the total amount of evidence necessary to trigger incongruent FAs exceeded the evidence underlying congruent FAs by a factor of 3.4, *p* < .001.

To summarize, just like Hits, FAs constitute a systematic reaction to foveal input. Congruency effects in FARs can therefore be interpreted in the same way as congruency effects in HRs: information about the peripheral saccade target interacts with congruent foveal input and facilitates its detection.

### Enhancement is spatially confined to the foveal region

The congruency effects described above may indeed reflect a spatially specific interaction between peripheral and foveal input. Alternatively, feature-based attention to the target’s orientation may yield a widespread enhancement of congruent orientations which could encompass the foveal region without being confined to it. To test if predictive enhancement is restricted to the pre-saccadic center of gaze and its immediate vicinity, we conducted a second experiment in which the probe, if presented, appeared in one of 37 locations spaced evenly on a horizontal axis from 4.5 dva to the left to 4.5 dva to the right of the screen center (in increments of 7-8 pixels). On each trial, the position of the probe was unpredictable for the observer. We determined congruent and incongruent HRs within a moving window including 6 adjacent locations and described the resulting spatial profiles by fitting Gaussian curves with position-invariant vertical offsets to individual observer data (**Figure 2D**; see ***Methods*** for details and ***Supplements*** for alternative fits). We flipped trials with leftward saccades, such that negative and positive position values indicate probe locations away and towards the saccade target, respectively. To account for small fixation errors, spatial profiles are aligned to the mean recorded fixation position during the saccade preparation period.

Across congruencies, HRs were highest in the center of gaze and decreased continuously as the eccentricity of the probe increased. While feature-based attention would predict a uniform detection advantage for congruent probes across all tested locations, the difference between the curves, and therefore enhancement, was most pronounced in the center of gaze (HR_cong-incong_ = 7.7±4.8% at −0.1±1.7 dva; *p* < .001). Enhancement remained significant within a region of 6.4 dva around the pre-saccadic fixation, extending further towards the saccade target than away from it: congruent HRs significantly exceeded incongruent ones from −2.6 to +3.8 dva. The congruent curve exhibited a significantly higher peak than the incongruent one (80.4±7.8% *vs* 87.9±7.3%, *p* < .001, *BF_10_* = 22.689). At the same time, and in opposition to a global enhancement of target-congruent orientations, the vertical offsets of the congruent and incongruent profile did not differ significantly (39.3±19.2% *vs* 35.9±16.3 %,*p* = .323, *BF_10_* = 0.340).

Enhancement may depend on the baseline performance level at a given eccentricity. In other words, sensitivity for peripheral probes may simply be too low for congruency effects to emerge. To ensure that an absence of enhancement for peripheral probes is a true consequence of their location, observers completed an additional session in which the probe appeared 3 dva to the left or the right of the screen center. Crucially, we adjusted the transparency of the probe against the background noise to elevate peripheral detection performances to a foveal level. While this manipulation was effective (**Figure 2D**), congruent and incongruent HRs differed neither for probes appearing away from (HR_cong-incong_ = −3.7±10.0%, *p* = .127, *BF_10_* = 0.531) nor towards (HR_cong-incong_ = 1.4±9.0%, *p* = .336, *BF_10_* = 0.352) the saccade target.

Again, congruent FARs significantly exceeded incongruent ones (FAR_cong-incong_ = 9.4±7.3%; *p* < .001). An inspection of noise properties revealed that – despite observers’ explicit knowledge about the possible range of probe locations – FAs were primarily triggered by foveal orientation information (see ***Supplements***).

### Post-saccadic target foveation boosts congruency

To investigate if post-saccadic target foveation influences pre-saccadic detection judgments, we removed the target during saccadic flight on half of the trials in Experiment 1 (*Pre*). On the remaining trials, the target remained visible for a brief duration after saccade offset (*Pre-Post; Mdn* = 22.3±4.12 ms). While we observed significant congruency effects when the target was visible exclusively during saccade preparation, congruency was more pronounced in *Pre-Post* trials: across all pre-saccadic time bins, the difference in HRs amounted to 2.5±6.0% in *Pre* and to 6.9±2.8% in *Pre-Post* trials, *p* < .001. Target foveation impacted the time course of congruency effects: when the target disappeared during the saccade, congruent HRs significantly exceeded incongruent ones in medium stages of saccade preparation ([–250 −200[ ms: HR_cong-incong_ = 4.4±4.9%; *p* = .003; [–200 −150[ ms: HR_cong-incong_ = 5.6±7.7%; *p* = .024; [–150 −100[ ms: HR_cong-incong_ = 8.4±7.6%; *p* < .001). Congruent and incongruent HRs in the earliest and latest time bins did not differ significantly (HR_cong-incong_ = −7.4±1.6%; 1.5±7.6%; *p*s = .060; .283). In *Pre-Post* trials, congruent HRs numerically exceeded incongruent HRs throughout saccade preparation, though this difference failed to reach significance in the earliest bin (HR_cong-incong_ = 5.6±7.5 %; *p* = .055). In all later bins, congruent HRs significantly exceeded incongruent HRs (HR_cong-incong_ = 7.6±8.3%, 8.2±8.3%, 3.8±5.7%, 10.5±9.5%; all *p*s < .041). A brief foveation of the target was sufficient to boost congruency. We determined the difference between congruent and incongruent HRs within a moving window (width: 10 ms; step size: 1 ms) sliding across the range of post-saccadic target durations (1-48 ms). Congruency was most pronounced in the window from 11 to 21 ms (HR_cong-incong_ = 9.1±3.4%). Congruent and incongruent HRs in the baseline bin were virtually identical for both *Pre* (HR_cong-incong_ = −0.9±9.8%, *p* = .534) and *Pre-Post* (HR_cong-incong_ = −0.4±12.6%, *p* = .590) trials.

## Discussion

We demonstrate that defining features of an eye movement target are predictively enhanced in the pre-saccadic center of gaze. Our findings reflect foveal enhancement rather than a general response bias to the target’s orientation, as the existence and magnitude of the congruency effect varied across spatial locations and time points.

### Is the fovea special? Potential mechanisms of foveal prediction

Enhancement was spatially specific, that is, most pronounced in the center of gaze and confined to an area of 6.4 dva around it. The true profile of enhancement may be even narrower: since our probe exhibited a diameter of 3 dva, congruency effects at eccentric locations may rely at least partly on the facilitated detection of the probe’s near-foveal margins. Could congruency effects with a similar spatial profile build up around any other relevant and therefore attended location in the visual field? The feasibility of peripheral congruency effects depends on the mechanism assumed to underlie our findings:

#### Joint modulation of spatial and feature-based attention

Foveal enhancement may rely on a joint modulation of spatial and feature-based attention^[26,27]^. Saccade preparation entails the emergence of two spatially confined attention pointers^[28]^: one centered on the saccade target and one centered on its predictively remapped location, i.e., the foveal region^[20,29]^. At the same time, the appearance of the saccade target in our investigation may have introduced global feature-based attention to its orientation^[30]^. The combination of spatial attention pointers carrying no feature information, and feature-based attention lacking spatial tuning may indeed achieve what we observe: a spatially specific alteration of visual sensitivity to defining features of a stimulus presented elsewhere in the visual field. It is conceivable that the fovea as the region of highest acuity is assigned a permanent attention pointer. In principle, however, this mechanism could operate across the visual field and may underlie previous findings that demonstrate an interaction of feature information between peripheral locations (for motion discrimination^[31]^, crowding^[32]^ and adaptation^[33–35]^).

Two spatially distinct, feature-selective attention pointers may account for the impact of post-saccadic target presence on the pre-saccadic time course of congruency effects in our investigation. Observers likely reported the presence of a target-congruent foveal probe whenever perceptual evidence for its orientation had exceeded a certain threshold. Assuming that orientation information was sampled simultaneously from the foveal and peripheral attentional focus, and assuming that the foci persisted for a brief period after saccade landing^[36,37]^, the salient post-saccadic foveal view of the target may have allowed even early and therefore subthreshold pre-saccadic foveal information to contribute to an above-threshold signal.

Nevertheless, this mechanism falls short of accounting for some aspects of our own as well as previous findings. First, we did not observe congruency effects when the probe appeared in one of two possible peripheral locations to which spatial attention pointers could have been allocated strategically. Second, congruency effects were aligned to the center of gaze rather than to the precise, predictively remapped location of the target, the coordinates of which depend on saccade endpoints on an individual-trial level^[29]^. Taking saccade endpoints into account reduced the spatial specificity of congruency effects (see ***Supplements***). Third, foveal retinotopic cortex contributes to the processing of complex peripheral shapes^[9–13]^. Whether feature-based attention can operate at this level of abstraction, and whether the proposed mechanism is viable for naturalistic objects involving feature conjunctions, is unclear. Consequently, the fovea may not merely be one of many locations an attention pointer can be allocated to. Instead, its unique characteristics seem to be harnessed in a way that would not be viable at other visual field locations.

#### Feedback connections to foveal neurons

Visual processing does not operate in a strictly feedforward fashion. For instance, neurons in several temporal areas (e.g., TE, IT, TPO, STS), most of which are associated with the computation of position-invariant object information, project back to primary visual cortex and potentially relay feature information about peripheral stimuli to V1 neurons with foveal receptive fields^[14, 15]^. Crucially, peripheral features can be decoded from brain activation in foveal – but not other peripheral – retinotopic areas^[9]^, supporting the fovea’s singular role as an active blackboard. Foveal feedback connections could account for all aforementioned findings: irrespective of the precise remapping vector, and despite the possibility to allocate peripheral attention pointers, feature information would be invariably relayed to the fovea. Moreover, temporal areas encode complex shapes, a coarse sketch of which could be fed back to foveal retinotopic cortex^[15]^. Average saccade metrics may still influence the profile of enhancement: while congruency effects in our study were most pronounced in the center of gaze, they extended further towards than away from the target. Since saccades are typically hypometric^[37,38]^ and the predictively remapped target location is therefore shifted towards the target by the magnitude of the undershoot^[28]^, an asymmetry in enhancement should have boosted congruency at the remapped target location across all trials. An involvement of foveal feedback connections in saccade preparation appears physiologically feasible: neurons in the aforementioned temporal areas exhibit median response latencies of 50-130 ms^[15]^. Feedback delays to foveal retinotopic cortex would thus lie well within the range of typical saccade latencies^[40]^. Indeed, our results are consistent with this timing: On trials in which the saccade target disappeared before saccade offset (*Pre*), we observed maximum enhancement when the foveal probe appeared 150-200 ms after the target (HR_cong-incong_ = 5.6±6.2%; *p* = .010).

### Is saccade preparation special?

While a link between foveal feedback and saccade preparation has been suggested repeatedly^[9,11]^, the influence of foveal processing on peripheral task performance has been studied almost exclusively during passive fixation. The only study investigating foveal feedback in an active setting revealed that a foveal distractor no longer impacted peripheral discrimination performance when observers prepared a saccade *away* from the to-be discriminated object^[10]^. These findings further support the main assumption motivating the current study: saccades automatically establish a transient and obligatory connection between the current and future foveal location, that is, the saccade target and the pre-saccadic center of gaze. A similar connection between foveal and peripheral input may exist or be inducible by task demands during fixation. Arguably though, this connection is strengthened considerably when a saccade successively projects two otherwise unrelated visual field locations (the current center of gaze and the saccade target) onto the same retinal location (the fovea). If foveal prediction relies on joint attentional modulations, congruency effects during fixation could mirror pre-saccadic ones only inasmuch as the dynamics of covert spatial attention resemble those of pre-saccadic attention and predictive remapping^[8]^. Irrespective of the mechanism underlying foveal prediction, interactions between target and fovea during saccade preparation are, at the very least, *accompanied by* simultaneous and obligatory attentional allocation to those locations. In accordance with this, we observed a pronounced increase in foveal HRs between the baseline and subsequent pre-saccadic timepoints (Experiment 1). Moreover, even though observers were aware that the probe could appear on an axis of 9 dva length, FAs were primarily triggered by foveal orientation information (Experiment 2; ***Supplements***). Both of these observations hold for target-congruent *and* incongruent orientations and suggest that during saccade preparation, visual attention was not only allocated to the saccade target but also to its future retinal location – the fovea.

### What is the function of foveal prediction?

Past investigations on foveal feedback required observers to make peripheral discrimination judgments. We, in contrast, did not ask observers to generate a perceptual judgment on the orientation of the saccade target. Instead, detecting the target was necessary to perform the oculomotor task. While the identification of local contrast changes would have sufficed to direct the eye movement, the orientation of the target enhanced target-congruent foveal processing. The automatic nature of foveal enhancement showcases that perceptual and oculomotor processing are tightly intertwined in active visual settings: planning an eye movement appears to prioritize the features of its target; commencing the processing of these features before the eye movement is executed may accelerate post-saccadic target identification and ultimately provide a head start for corrective gaze behavior^[16–18]^. Since we routinely direct our gaze to relevant information rather than inspecting it peripherally, the foveation of peripheral input via an eye movement may not merely be another instance of foveal feedback, but the very reason for its existence.

## Conclusion

We suggest a crucial contribution of foveal processing to transsaccadic visual continuity which, up until now, has been overlooked. Feedback connections to foveal retinotopic cortex, so far characterized during passive fixation, gain a predictive nature during saccade preparation. They entail a retinotopic anticipation of soon-to-be foveated information – notably without any need for predictive spatial updating. As a behavioral consequence, the predictive foveal enhancement of target features demonstrated here may contribute to the continuous perception of eye movement targets and accelerate post-saccadic gaze correction.

## Acknowledgments

We thank Jude Mitchell and Shanna Coop for helpful discussions on foveal prediction, Lea Krätzig for help with data collection, and the members of the Active Perception and Cognition group for participating in our experiments in times of need.

## Methods

### Experiment 1

#### Participants

Seven human observers (five females, seven right-handed, four right-eye dominant, one author) aged 22 to 34 years (*Mdn* = 28.0) participated in the experiment. Since we could not derive effect-size estimations from prior studies, we chose a sample size within the typical range of experiments investigating pre-saccadic attention shifts^[41,22]^. Normal (*n* = 4) or corrected-to normal (*n* = 3) visual acuity was ensured at the beginning of the first session using a Snellen chart (Hetherington, 1954) embedded in a Polatest vision testing instrument (Zeiss, Oberkochen, Germany). Observers yielding scores of 20/25 or 20/20 were invited to proceed with the experiment. Ocular dominance was assessed using the Miles test^[42]^. Since data collection was carried out during the COVID-19 pandemic, our sample was composed of either lab members (*n* = 6) or external participants recruited through word of mouth (*n* = 1). Apart from the author, all observers were naïve as to the purpose of the study. Participants gave written informed consent before the experiment and were compensated with either accreditation of work hours, course credit or a payment of 8.50€/hour. In case of monetary compensation, a bonus of 6€ was added upon completion of all seven sessions. The study complied with the Declaration of Helsinki in its latest version and was approved by the Ethics Committee of the Department of Psychology at Humboldt-Universität zu Berlin. The research question, experimental paradigm and data analyses were preregistered on the Open Science Framework (https://osf.io/xr2jk). Raw and pre-processed data are publicly available at *[link will be made available upon acceptance]*.

#### Apparatus

##### External setup

The experiment was conducted in a dimly-lit, sound-dampened booth. Stimuli were projected on a 200 × 113 cm screen (Celexon HomeCinema, Tharston, Norwich, UK) using a PROPixx DLP Projector (Vpixx Technologies, Saint-Bruno, QC, Canada) with a spatial resolution of 1920 × 1080 pixels and a refresh rate of 120 Hz (frame duration of 8.3 ms). Observers faced the screen at a viewing distance of 180 cm while their heads were stabilized on a combined chin and forehead rest. Throughout the experiment, the position of the dominant eye was recorded at a sampling rate of 1 kHz using a desk-mounted infrared eyetracker (EyeLink 1000 Plus; SR Research, Osgoode, Canada). Stimulus presentation was controlled by a DELL Precision T7810 Workstation (Debian GNU Linux 8) and implemented in Matlab 2016b (Mathworks, Natick, MA, USA) with the PsychToolbox^[43,44]^ and Eyelink toolbox^[45]^ extensions. Observers generated their responses on a standard QWERTY keyboard positioned centrally in front of them.

##### Stimulus generation

To generate the background noise images, we applied a fast Fourier transform to uniform white noise, multiplied the noise spectrum with its inverse radial frequency, and transformed it back using an inverse fast Fourier transform^[21,22]^. A total of 34 noise images (17 for the staircase block; 17 for the main experiment) were generated for each observer and session before the respective task. While each noise image appeared in every trial, the order of presentation was randomized across trials. Within a trial, a certain noise image was never repeated. Assuming that nine noise images were presented between the onset of the noise stream and the onset of the saccade, and disregarding the order of presentation (see ‘Data Analysis’), a total of 24,310 image combinations (i.e., 17!/9!(17 – 9)!) were possible in the critical time window. Across all 833 images presented in the main experiment (7 observers x 7 sessions x 17 images), the power spectral density (PSD) decreased by 5.07 dB per octave (see **Figure 1B** for the PSD of all noise images presented in a randomly chosen session).

To create the probe and target patches, we determined the background noise image that would be on screen at the desired probe presentation time and selected the pixel values in the relevant spatial region (3×3 dva) of that image. We subsequently filtered the orientation content of the selected noise window to −45° or 45° in the main experiment (**Figure 4 in *Methods***, ‘Stimulus generation’), and to −45°, 0° or 45° in the staircase block. The width of the orientation filter representing the sharpness of tuning around the desired orientation was constant (α_FiltWidth_ = 20°). We adjusted the difficulty of the foveal detection task by varying the transparency of the probe against the noise background. To guarantee a smooth transition between the noise background and overlayed patches, we superimposed all patches with a raised 2D cosine mask (radius = 1.5 dva; sigma = 0.7 dva^[21]^). Note that, whereas the probe appeared for the duration of a single noise image, the presentation of the saccade target spanned multiple images. Since the target constituted an orientation-filtered version of the respective background region, the appearance of the target changed dynamically in 50 ms intervals.

**Figure 4.**
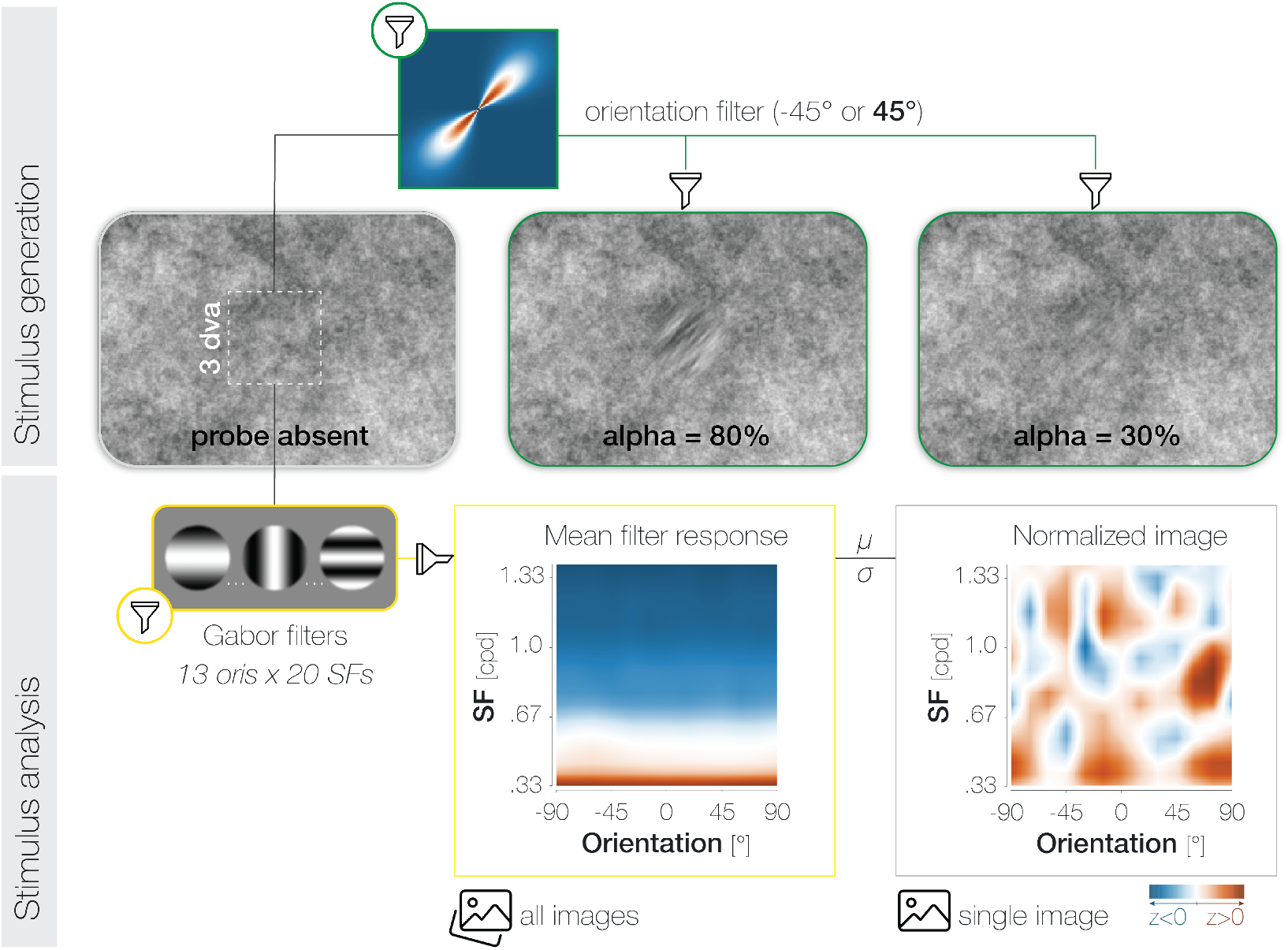
Stimulus generation (top) and analysis (bottom). To generate probe and target patches, we applied smooth orientation filters (green lines) to the relevant regions of the respective noise image. The 2D Fourier transform of a 45° filter is plotted for illustration. To adjust task difficulty for individual observers and different experimental purposes, we varied the opacity a (1-transparency) of the probe. High α-values increase visibility (see α = 80%) while low α-values decrease visibility (see α = 30%). Note that the calibration of the displaying device (in our case the projector) and the constant stimulus flicker in our experiment influence the visibility of the probe stimulus and should be considered when judging the displayed examples. To analyze the properties of the foveal noise window on probe absent trials, we computed its dot product with 260 Gabor filters with varying orientation (ori)*spatial frequency (SF) characteristics (yellow outlines). The resulting mean filter response across all images shows high energy for low-SF information (orange) and low energy for higher SFs (blue). To account for the asymmetry in power across SFs, we normalized every image by the mean (μ) and standard deviation (σ) of the set of images presented in a given experimental session for a given observer.

#### Procedure

Every participant completed seven sessions within a mean span of 11 days (min = 7 days; max = 17 days). Individual sessions were performed on several days and lasted approximately 90 min. Each session started with a staircase block, followed by the main experiment. To familiarize observers with the task, the staircase procedure was preceded by a slow-motion training block and an eye movement training block in the first session. Both training blocks were continued until reliable task and oculomotor performance was achieved. In all parts of the experiment, observers monitored the onset of a peripheral saccade target and, upon its detection, prepared an eye movement towards it. In 50% of all trials, and at different time points before the eye movement, a probe stimulus appeared in the screen center, i.e., observers’ current, pre-saccadic center of gaze. Observers executed the eye movement and subsequently reported if the probe in the screen center had been present or absent. In case of a ‘present’ response, they additionally reported the perceived orientation of the foveal probe (2-AFC: left vs right).

##### Trial procedure in the main experiment

At the beginning of each trial, a white fixation dot of diameter 0.3 dva was presented in the center of a static noise image covering the entire screen (**Figure 1A**). After stable fixation had been determined within a circle of 2.0 dva radius around the dot for at least 200 ms, the first noise image remained visible for a random duration of 550 to 1050 ms. Afterwards, the background noise started flickering at a temporal frequency of 20 Hz, corresponding to a presentation duration of 50 ms for each image. Following a delay of 200 ms after noise stream onset, the saccade target, i.e., an orientation-filtered patch with a diameter of 3 dva, appeared 10 dva to either the left or the right of fixation. Observers were instructed to move their eyes to the target as fast as possible. The saccade target was oriented either −45° (corresponding to a leftwards tilt) or +45° (corresponding to a rightwards tilt) and appeared at an opacity of *a* = 60% against the background noise. Since no additional saccade cue was presented, the detection of the target was necessary to perform the oculomotor task.

On 50% of trials, the probe, i.e., a second orientation-filtered patch of 3 dva diameter, appeared in the screen center. Crucially, the probe’s orientation was either −45° (leftwards tilt) or +45° (rightwards tilt) and therefore either congruent or incongruent to the target. If present, the probe remained on screen for the duration of one noise image (50 ms). To obtain estimates of visual sensitivity throughout the saccade preparation period, we defined three possible delays between target and probe onset. The shortest delay did not vary across observers: the probe was presented 50 ms after target onset. To ensure that the probe disappeared shortly before or at saccade onset on a sufficient number of trials, the longest delay was set to an observer’s median saccade latency in the preceding staircase block minus the duration of the probe (50 ms). The intermediate delay was chosen to fall right between the shortest and longest one. We slightly adjusted all delays, such that the on- and offset of the probe always coincided with the on- and offset of one background noise image in the stream. This timing protocol allowed us to obtain sensitivity measures throughout the saccade preparation period while minimizing offline trial loss due to intra- and post-saccadic probe presentations. To obtain a baseline performance estimate outside the saccade preparation period, we added one further onset condition in which the probe appeared 150 ms *before* the saccade target.

We investigated if foveating the saccade target for a brief period of time after saccade landing would influence pre-saccadic detection judgements. For this purpose, the experiment involved two trial types: on half of the trials (*Pre*), the saccade target was removed from screen as soon as recorded gaze position left a circle of 2 dva radius around the fixation dot. On the other half (*Pre-Post*), the saccade target remained visible for a short duration after gaze position had entered a circle of 3.25 dva radius around the center of the saccade target (see ‘Gaze-contingent timing and online eye movement criteria’ for details). Across all sessions and observers, the saccade target was foveated for *Mdn* = 22.6±4.12 ms after saccade offset (determined offline; **Figure 3B**).

Irrespective of probe and target timing, the background images flickered for a total duration of 800 ms. Afterwards, the last image remained on screen throughout the response period. Observers indicated if the probe in the screen center had been present or absent by pressing the up- or downarrow key on a standard keyboard in front of them. After a ‘present’ response, they additionally reported the perceived orientation of the probe by pressing the left or right arrow key. We instructed observers to prioritize the presence/absence judgment over the orientation judgment. Specifically, they were encouraged to guess on the second response rather than responding ‘absent’ if they perceived the probe stimulus but were unsure about its orientation. The next trial was initiated after an intertrial interval of 500 ms during which the last noise image in the stream remained on screen.

##### Trial procedure in the staircase block

Before the main experiment, we administered a staircase block to adjust the α-level, i.e., the opacity (1–transparency) of the foveal probe against the background noise, to an optimal level. The trial procedure was identical to the main experiment with the following exceptions: First, and quite obviously, the α-level of the probe was adjusted adaptively in the staircase block but remained constant within each session of the main experiment. Second, to adjust the α-level in the absence of potential congruency effects, and to avoid rendering observers more familiar with the subset of incongruent trials before the main experiment, the saccade target was oriented vertically (0°) on all staircase trials. The probe stimulus was tilted 45° to the left or right, mirroring the main experiment. Third, while the probe was presented at one of four time points in the main experiment, it appeared 50 ms after target onset on all staircase trials. We implemented this measure to avoid ceiling performance for incongruent probes in early stages of saccade preparation when we assumed foveal sensitivity to be highest. Lastly, the target was removed during the eye movement on all staircase trials.

##### Session structure

###### Task familiarization (session 1)

In the first session, we familiarized observers with the task by presenting a random subset of trials in a slowed down version and in the absence of oculomotor requirements. For this purpose, stimulus presentation times were increased by a factor of six. Participants generated verbal replies on the location of the saccade target, the presence or absence of the foveal probe, and its perceived orientation after a ‘present’ judgment. Once an observer was able to perform the task at the current speed, presentation times were gradually reduced until reliable task performance was achieved at normal speed. Observers subsequently performed eye movement practice trials until comfortable with the oculomotor aspect of the task.

###### Staircase procedure (all sessions)

The α-level of the foveal probe was adjusted following a single-interval adjustment matrix protocol (SIAM^[46]^). For each observer and session, we aimed to estimate the α-level at which a maximum reduced HR (HR–FAR) of 0.5 would be obtained. Initially, the probe was presented at an opacity of 50%. Possible α-levels ranged from 12% to 100%, in 2% increments. The α-level was adjusted after each trial based on the type of response generated: After Hits, it was reduced by 12%. After Misses and FAs, it was increased by 12% and 24%, respectively. No contrast adjustment was administered after Correct Rejections. The orientation report following a ‘present’ response did not affect the adjustment of α-levels. Initial step sizes were halved after the first and second reversal. Step sizes were reset if five consecutive Hits were generated at the same α-level. The staircase block terminated after 96 completed trials and took observers approximately 10 minutes to complete. The resulting opacity estimate was obtained by averaging α-values corresponding to the last six reversals. If fewer than six reversals had occurred, all available reversals were averaged to obtain the α-estimate.

###### Main experiment (all sessions)

In each session, the main experiment was divided into six blocks of 107 trials each. Breaks were offered after every 54 trials. After the first block half, an observer’s current HR and FAR along with the resulting d’ score across all conditions was displayed on screen. Performance measures were not labelled or otherwise interpretable for the observer. Prior to data collection, we had specified α-level adjustments in case of exceedingly good or poor performance (see preregistration). In the course of data collection, we realized that, with increasing session number, observers tended to generate more FAs in the staircase block. This was likely the combined effect of repeated exposure to probes with high transparency and a training-induced increase in sensitivity to weak orientation content in the background noise. In general, we argue that FAs in our investigation may reflect an enhancement of orientation information rather than purely unsystematic response behavior that would necessitate a decrease in task difficulty (see ‘Results’). To avoid a saturation of HRs in the main experiment, and to reduce the number of iterations needed to reach the targeted performance range, we therefore reduced the α-level by more than the preregistered step size (5%) at a time if considered appropriate. Across all observers and sessions, the probe was presented at a median opacity of α = 25.0±2.9% (median per session: 31.4%, 24.0%, 25.0%, 25.0%, 25.0%, 25.0%, 22.0%). The main experiment took observers around 60 minutes to complete.

##### Gaze-contingent timing and online eye movement criteria

In *Pre-Post* trials, we intended to time the disappearance of the saccade target such that it would remain visible for 16–24 ms (i.e., 2 to 3 refresh frames) after eye movement landing. Achieving this required an estimate of the time interval between the boundary cross and the true offset of the saccade determined after the testing session. As an initial estimation in the first session, we presented the target for 3 frames after the recorded gaze position first crossed a boundary of 3.25 dva around the target. After every session, we inspected the post-saccadic presentation duration in offline analyses and adjusted the number of target frames accordingly in the following session. Trials were assigned to the *Pre* or *Pre-Post* condition based on an offline inspection of saccade and stimulus timing (**Figure 3B**): trials in which the target was supposed to disappear intra-saccadically but was still visible after the eye movement (mean *n* = 9 trials per observer) were assigned to the *Pre-Post* condition. Trials in which the target was supposed to be visible after the eye movement but disappeared during saccadic flight were assigned to the *Pre* condition (mean *n* = 42 per observer).

A trial was aborted if at any time before target onset, gaze position was recorded further than 2.0 dva away from the pre-saccadic fixation dot. After target onset, gaze position had to cross this 2.0-dva threshold within 400 ms, in the direction of the target. Saccade landing had to be recorded within a circle of radius 3.0 dva around the center of the target. Note that target offset was time-locked to a slightly larger boundary (3.25 dva) which had achieved most accurate timing during piloting. To control post-saccadic foveal input, observers’ gaze position had to remain within a circle of radius 5.5 dva around the target until the first keyboard response had been generated. Error-specific feedback messages were displayed after trial abortions. Aborted trials were appended at the end of a given block.

#### Data analysis

##### Eye movement pre-processing

The pre-processing of eye movement data and all subsequent analyses were implemented in Matlab 2018b (Mathworks, Natick, MA, USA). Saccades were detected offline using a velocity-based saccade detection algorithm^[47]^. We defined saccade onset as the time point at which the current eye velocity had exceeded the median eye velocity from all preceding samples by 5 SDs for at least 8 ms. When recorded with pupil-based eye trackers, saccades often exhibit post-saccadic oscillations, which are assumed to reflect residual pupil movement rather than a true rotation of the eyeball^[48]^. We effectively excluded them from the estimated saccadic profile by not merging detected saccadic events separated by one sample or more.

We excluded trials in offline analyses if no response saccade had been generated in the critical time window between 150 ms before and 550 ms after cue onset, if a saccadic event was detected before the actual response saccade and/or if gaze position samples were missing at any time before saccade onset. Moreover, trials involving anticipatory response saccades, i.e., saccades with a latency below 80 ms, were excluded. Finally, to ensure that post-saccadic foveal input was stable on the retina for a certain period of time after saccade landing, we excluded trials in which a second saccade had occurred between 25 and 100 ms after response saccade offset (‘crit’ in **Figure 3B**). Since post-saccadic oscillations were often registered as a second saccadic event, we introduced a short time window of 25 ms after response saccade offset, during which saccadic activity did not lead to trial exclusions. Combined, these criteria resulted in an offline exclusion of 8.12% of trials. We excluded a further 4.62% of trials in which the foveal probe disappeared intra- or post-saccadically rather than before saccade onset. A total of 27,328 trials were carried on to further analyses. The parameters of all included response saccades are given in the ***Supplements***.

##### Analysis of noise content

We inspected the foveal noise content in probe absent trials separately for congruent FAs (observers reported perceiving the target’s orientation) and incongruent FAs (observers reported perceiving the non-target orientation). On each trial, we determined the noise images that had been visible for their full duration during the entire potential probe presentation period, i.e., from the onset of the dynamic noise stream to the onset of the saccade (mean *n* = 9.2). To isolate the impact of saccade preparation, we subsequently separated images that had preceded the onset of the saccade target (baseline; *n* = 4) and images visible from the onset of the saccade target to saccade onset (saccade preparation; mean *n* = 5.2; **Figure 3A**).

To investigate the relation between response behavior and noise content at the potential probe location, we selected all pixels within a square of 3 dva side length around the screen center. We described the visual properties of each noise window along two dimensions: its spatial frequency (SF) and orientation (ori) content. To determine the energy of a certain SF*ori combination in the noise window, we created two Gabor filters with the corresponding properties (3 dva in diameter; one in sine and one in cosine phase^[49–51]^). We then convolved the pixel content in the noise window with both filters and averaged their responses. Note that Gabor filters with orientations from −π/2 to π/2 will yield slightly different responses than those with orientations from −π/2+π to π/2+π even though they effectively correspond to the same orientation^[51]^. We accounted for this by applying filters for orientations from −π/2 to π/2 as well as their counterparts from −π/2+π to π/2+π, and averaging their responses^[50]^.

Using this method, we obtained filter responses for 260 SF*ori combinations per noise image (**Figure 4 in *Methods*, ‘Stimulus analysis’**). SFs ranged from 0.33 to 1.39 cpd (in 20 equal increments). Orientations ranged from −90 to 90° (in 13 equal increments). To normalize the resulting energy maps, we z-transformed all filter responses using the mean and standard deviation of filter responses from the set of images presented in a certain session. To obtain more fine-grained maps, we applied 2D linear interpolations by iteratively halving the interval between adjacent values 4 times in each dimension. To facilitate interpretability, we flipped the energy maps of trials in which the target was oriented to the left. In all analyses and plots, +45° thus corresponds to the target’s orientation while −45° corresponds to the other potential probe orientation. Noise images and analysis scripts can be found on *[link will be made available upon acceptance]*.

##### Tests of statistical significance

We used bootstrapping to compare HRs and FARs between time points and congruency conditions. Within each observer, we determined the means in the to-be compared conditions and computed the difference between those means. Across observers, we drew 10.000 random samples from these differences (with replacement). Reported *p*-values correspond to the proportion of differences smaller than or equal to zero. We considered *p*-values <= 0.05 significant.

We used a two-step approach to identify SF*ori combinations that exhibited a significantly high or low energy in the foveal noise region when a congruent or incongruent FA was generated. First, we contrasted the energy of every SF*ori combination to zero (i.e, to the mean of the standard normal distribution for this specific combination), using two-sided one-sample t-tests. We then selected SF*ori combinations with a *t*-value > 1.94 (corresponding to a significance level of *p* = 0.05) and grouped neighboring above-threshold combinations into a cluster using the Matlab function bwconncomp (pixel connectivity = 4).

Two orientations were behaviorally relevant in our experiment: −45° and +45°. To constrain our analyses, we therefore selected the two clusters with the highest summed *t*-value per response type and carried only those on to further tests. Indeed, all clusters identified in the first step were lateralized, i.e., exhibited centers of mass close to −45° or +45°. Since we had not corrected for multiple comparisons when identifying the clusters, we verified their effective meaningfulness using a non-parametric bootstrapping approach. Specifically, we tested whether filter energies within a cluster differed significantly from their SF-matched equivalents around the other relevant orientation. For this purpose, we determined the sum of filter responses within each cluster on an individual-observer level. To provide a test of lateralization, we flipped individual-observer maps in the orientation dimension – such that −45° now corresponded to +45° and vice versa – and summed the flipped filter responses within the non-flipped clusters. We then contrasted the filter responses for original and flipped maps by drawing 10,000 bootstrapping samples from our set of observers (with replacement) and computing the mean difference in filter responses for each sample. A cluster was reported in the ‘Results’ section if ≤ 5% of bootstrapped samples yielded a difference between original and flipped maps that was larger than the true difference. Reported *p*-values indicate this proportion. Three out of four clusters identified in the first step proved lateralized and are reported in the main text.

We would like to point out that, just like cluster-based permutation tests^[53,54]^, our approach does not establish the significance of single points within a cluster. While we can conclude that filter responses in a reported cluster differ significantly from filter responses in the same region of orientation-flipped maps, we cannot conclude that this only holds for a cluster with the exact dimensions reported. In consequence, and identically to cluster-based permutations, statements on the precise extent of a cluster along the orientation and SF axes are descriptive rather than inferential in nature.

### Experiment 2: Spatial specificity

To test if predictive enhancement is confined to the pre-saccadic center of gaze, we presented the probe stimulus not only in the screen center but on a horizontal axis from 4.5 dva to the left to 4.5 dva to the right of the screen center (sessions 1-6). In session 7, we intended to ensure that a potential absence of enhancement for peripheral probes is a true consequence of probe location and cannot be attributed to differences in baseline sensitivity. For this purpose, the probe was presented at one of two possible locations – either 3 dva to the left or 3 dva to the right of the screen center – and appeared at an α-value adjusted to approximate foveal performance levels. The remaining experimental procedure remained largely unaltered. Differences to Experiment 1 are explained below.

#### Participants

Nine human observers (seven females, all right-handed, seven right-eye dominant; no authors) aged 22 to 34 years (*Mdn* = 28.9) participated in the experiment. Six observers had participated in Experiment 1. We chose to increase our sample size since we removed the target during the eye movement on all trials of Experiment 2, a condition (*Pre*) in which we observed smaller congruency effects than in the *Pre-Post* condition in Experiment 1 (across all pre-saccadic time bins: *HR_cong-incong_* = 2.52% vs 6.85%). Normal (*n* = 5) or corrected-to normal (*n* = 4) visual acuity was ensured at the beginning of the first session using a Snellen chart (Hetherington, 1954) embedded in a Polatest vision testing instrument (Zeiss, Oberkochen, Germany). Again, as data collection was carried out during the COVID-19 pandemic, our sample was composed of either lab members (*n* = 8) or external participants recruited through word of mouth (*n* = 1). Consent procedures, participant compensation and ethics approvals were identical to Experiment 1. The experimental question, paradigm and analyses were preregistered on the Open Science Framework (https://osf.io/6s24m). Raw and pre-processed data are publicly available at *[link will be made available upon acceptance]*.

#### Apparatus and stimulus properties

The external setup was identical to Experiment 1. To allow for a larger variation of foveal noise content across trials, we increased the number of noise images generated per session from 34 to 70 (35 for the staircase block, 35 for the main experiment). In consequence, only a random subset of all generated noise images appeared on a single trial (in randomized order). Again, a certain noise image was never repeated within a trial. Assuming that nine noise images were presented between the onset of the noise stream and the onset of the saccade, and disregarding the order of presentation, a total of 70,607,460 image combinations (i.e., 35!/9!(35 – 9)!) were possible in the critical time window. Across all 2205 images presented in the main experiment (9 observers x 7 sessions x 35 images), the power spectral density (PSD) decreased by 4.89 dB per octave.

Target and probe patches were generated identically to Experiment 1 with the exception that, across trials, probe patches constituted a filtered version of the background noise not only in the screen center but at all possible probe locations.

#### Procedure

Every participant completed seven sessions within an average span of 18 days (min = 15 days; max = 28 days). Individual sessions were performed on several days and lasted approximately 90 min. Like in Experiment 1, each session started with a staircase block, followed by the main experiment. If an observer had not participated in Experiment 1, the staircase procedure was preceded by a slow-motion training block and an eye movement training block in the first session (see Experiment 1).

##### Staircase block

###### Sessions 1-6

The trial procedure was identical to the staircase block in Experiment 1: observers executed a saccade to a vertically oriented target that disappeared during the eye movement and detected the presence or absence of a probe stimulus in the screen center. No peripheral probes were presented – the contrast adjustment targeted an optimal performance level for the presence/absence judgment of foveal probes. Since we often had to reduce the estimated α-value to avoid ceiling performance in Experiment 1, we modified the classical SIAM protocol for our purposes in Experiment 2:

Initially, the foveal probe was presented at an α-level of 30%. Possible values ranged from 14% to 100%, in 2% increments. After each trial, the probe’s *a* was adjusted depending on the type of response generated: After Hits, it was reduced by 18%. We did not increase *a* more after an FA than after a Miss, deviating from the classical SIAM protocol: after Misses and FAs, α was increased by 27%. We implemented this measure because FAs in our design can indicate *increased* sensitivity to target-congruent orientation information in the foveal noise region. As the SIAM staircase assumes changes in FARs to result from changes in decision criteria rather than sensitivity to external signals, this property had likely inflated α-values in the staircase block of Experiment 1.

###### Session 7

In the last session, we aimed to determine the α-level at which observers would yield the same incongruent HR for probes presented at 3 dva eccentricity as they did for a foveal probe in the preceding sessions. For this purpose, we randomly interleaved trials in which the probe was presented 3 dva to the left and 3 dva to the right of the screen center, rendering the location of the probe unpredictable on an individual-trial level. After Hits, the current α-value was reduced by 18%. After Misses and FAs, it was increased by 14.7%. No α-adjustment was administered after Correct Rejections.

In all seven sessions, initial step sizes were halved after the first and second reversal. Step sizes were reset if five sequential hits had been generated at the same α-level. The staircase terminated after 96 trials and took observers 10-15 minutes to complete. The resulting α-estimate was obtained by averaging α-values corresponding to the last eight reversals. If fewer than eight reversals had occurred, all reversals were averaged to obtain the α-estimate.

##### Main experiment

###### Sessions 1-6

The trial procedure in sessions 1-6 was identical to the procedure in Experiment 1 with the following exceptions: First and foremost, the probe could appear in one out of 37 potential locations. These locations were spaced evenly on a horizontal line extending from 4.51 dva to the left to 4.51 dva to the right of the screen center, in increments of 0.232 or 0.265 dva (i.e., 7 or 8 pixels). Different probe locations were randomly interleaved. Second, we intended to isolate purely pre-saccadic influences of saccade target features on foveal sensitivities and removed the saccade target during the eye movement on all trials. Based on offline analyses, the target disappeared *Mdn* = 15±4.30 ms before saccade offset. Lastly, to obtain reliable spatial profiles within a feasible number of experimental sessions, we chose to present the probe at a single, optimal pre-saccadic time point, i.e., 100 to 75 ms before the eye movement. In this bin, congruency effects had been most pronounced in the *Pre* condition of Experiment 1 (see **Figure 3B**). Again, we relied on the median saccade latency recorded in the staircase block of each session to time the onset of the probe in the main experiment. Based on offline analyses, the probe disappeared *Mdn* = 96.28±18.13 ms before the eye movement.

The main experiment involved 8 blocks of 74 trials each. Breaks were offered between blocks. After the first block, the observer’s current HR and FAR along with the resulting d’ score across all probe positions and conditions was displayed on screen (not labelled or otherwise interpretable for the observer). If the difference between the HR and the FAR was below 15%, the probe’s a was increased by 5%. If the difference between HR and FAR was above 40%, the probe’s a was decreased by 5%. In Experiment 1, we experienced that an observer’s subjective report on task difficulty, paired with an inspection of their performance after each experimental session, was a reliable indicator of targeted performance levels. We therefore preregistered the option of deviating from this adjustment routine to be able to correct the staircase contrast in fewer iterations, and resorted to this option if necessary. Across observers and sessions, the probe was presented at a median opacity of α = 28.3±7.6% (median per session: 33.0%, 28.9%, 29.6%, 27.8%, 25.2%, 28.2%). The main experiment took observers around 60 minutes to complete.

###### Session 7

The main experiment in the last session was identical to the main experiment of previous sessions with one exception: the probe appeared either 3 dva to the left or 3 dva to the right of the screen center. Observers were explicitly informed about the placement of the probe. Trials of both conditions were randomly interleaved.

Every observer completed 9 blocks of 64 trials each. Breaks were offered between blocks. After the first block, we adjusted the probe’s a in case of exceedingly good or poor performance similarly to previous sessions. Across observers, the probe was presented at a median opacity of α = 39.0±11.1%. The main experiment took observers around 60 minutes to complete.

###### Gaze-contingent timing and online eye movement criteria

During the experiment, we removed the saccade target once gaze had crossed a boundary of 2.0 dva around the pre-saccadic fixation. Trial abortion criteria were identical to Experiment 1.

#### Data analysis

##### Eye movement pre-processing

Pre-processing parameters and exclusion criteria were identical to those in Experiment 1. We excluded 11.25% of trials due to saccade characteristics and a further 2.29% of trials due to intra- or post-saccadic probe presentations. We furthermore excluded all trials in which the target had been visible after saccade landing (0.89% of trials). On all included trials, the target disappeared before saccade offset (sessions 1-6: *Mdn* = −15.00±4.30 ms; session 7: *Mdn* = −15.11±5.33 ms). A total of 31,674 trials were carried on to further analyses. The parameters of all included response saccades are given in the ***Supplements***. While saccade latencies and amplitudes showed small-scale variations across probe locations, these variations did not influence the spatial profile of enhancement.

##### Function fitting

We determined congruent and incongruent HRs within a moving window including 6 adjacent locations at each iteration (i.e., 1.46 dva; step size = 0.03 dva) and fitted Gaussian functions with constant vertical offsets to the resulting spatial profiles. Before the fitting step, we realigned all probe locations to the mean fixation position recorded during the saccade preparation period from target onset to saccade onset. We furthermore flipped probe locations in trials with leftwards saccades. In consequence, an x-axis value of zero indicates that the probe was presented in the center of gaze. Negative and positive x-values denote that the probe appeared in the opposite (‘away’) or same (‘towards’) hemifield of the saccade target, respectively. All position values indicate horizontal coordinates. We related HRs (y-axis) to probe locations (x-axis) using a Gaussian distribution with a constant offset from zero:

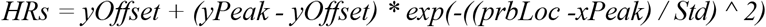

where *yOffset* refers to the constant vertical offset of the Gaussian curve, *yPeak* and *xPeak* refer to the vertical and horizontal coordinate of its peak, respectively, and *Std* refers to its standard deviation. Fitting was performed on an individual-observer level using a nonlinear least-squares fitting protocol (Matlab function ‘lsqnonlin’; trust-region-reflective algorithm). We defined lower and upper bounds of 0 and 0.7 for *yOffset*, 0.3 and 1.0 for *yPeak*, 1 and 31 for *xPeak* and 0 and 31 for *Std*. To account for the reliability of each data point in the fitting process, we minimized the deviation between measured values and a weighted cost function. We obtained the weighted cost function by determining the deviations between observed HRs and HRs predicted with the current set of parameters at each iteration. We subsequently multiplied these deviations with a weight vector directly proportional to the number of trials contributing to a given data point. Across all observers and data points, assigned weights ranged from 0.62 to 1 for congruent HRs and from 0.63 to 1 for incongruent HRs. The minimum and maximum number of trials in an individual-observer data point amounted to 34 and 126, respectively. To obtain mean HR profiles across observers for plots (**Figures 2C,S1,S3**), we determined a Gaussian curve with each observer’s individual parameter estimates and averaged those curves.

The fitted profiles closely approximated observed HRs, with a mean absolute error of 0.5% for congruent and 1.1% for incongruent HRs. However, while raw and fitted spatial profiles exhibited a slightly asymmetric shape across observers, the fitted Gaussian curves were strictly symmetrical on an individual-observer level. The asymmetric shape of the mean fitted profile likely relies on a variation of the *xPeak* parameter or a joint variation of the *xPeak* and *Std* parameter on an individual-observer level. To allow for asymmetric profiles within each observer, we fitted two-term Gaussian functions to individual-observer data (see ***Supplements***). Since both approaches provided equivalent fits, and since the Gaussian curve with vertical offset involves fewer yet more easily interpretable parameters, we decided to present these fits in the ***Results*** section. Nonetheless, fitting two-term Gaussian functions to our data did not alter the nature of findings: enhancement was highest in the center of gaze and significant within a similar spatial range. All statistical comparisons reported in the ***Results*** section rely on bootstrapping analyses (with replacement; *n* = 10.000; see Experiment 1). Whenever we relied on a null effect to support a specific claim (e.g., the spatial specificity of enhancement), we supplemented *p*-values with Bayes Factors to gage evidence for the absence of an effect (*BF_10_s*, scale factor 0.707). Bayes Factor calculations rely on two-sided, one-sample t-tests. Only the test contrasting the peak parameters of the fitted Gaussian profiles was directional (congruent > incongruent) since both potential mechanisms – spatially specific and global enhancement – predict this pattern.

## Supplementary Materials

### Eye movement parameters in Experiment 1

On average, observers executed hypometric saccades with a median amplitude of 9.30±.57 dva for leftwards and 9.74±.72) dva for rightwards saccades, and yielded a median saccade latency of 279.14±16.01 ms (leftwards saccades: *Mdn* = 277.57±23.19 ms; rightwards saccades: *Mdn* = 279.21±12.60 ms). Saccade latencies were stable across stimulus and response conditions, suggesting that the presentation of the foveal probe did not alter eye movement preparation: we observed comparable latencies for probe present and probe absent trials (279.57 ms *vs* 278.57 ms, *t*(6) = 1.32,*p* = 0.234), Hits, Misses, FAs and CRs (281.14 *vs* 275.29 *vs* 273.79 *vs* 280.71 ms, *F*(3,24) = 0.39,*p* = 0.761, one-way ANOVA), Hit trials with target-congruent and target-incongruent foveal probes (284.29 *vs* 282.71 ms, *t*(6) = 1.72, *p* = 0.052), and FA trials with target-congruent and target-incongruent orientation reports (273.64 *vs* 273.00 ms, *t*(6) = 0.34,*p* = 0.748).

### Eye movement parameters in Experiment 2

Observers executed hypometric saccades with a median amplitude of 9.37±1.09 dva for leftwards and 9.70±.96 dva for rightwards saccades. The median saccade latency was 271.50±26.40 ms (leftwards saccades: *Mdn* = 273.00±30.42 ms; rightwards saccades: *Mdn* = 271.50±22.15 ms). Again, latencies were stable across stimulus and response conditions: we observed comparable latencies for probe present and probe absent trials (266.78 *vs* 269.33 ms, *t*(8) = −1.90, *p* = 0.094), Hits, Misses, FAs and CRs (266.67 *vs* 267.06.29 *vs* 268.79 *vs* 269.79 ms, *F*(3,32) = 0.03,*p* = 0.994, oneway ANOVA), Hit trials with target-congruent and target-incongruent foveal probes (266.56 *vs* 267.11 ms, *t*(8) = −0.351, *p* = 0.735), and FA trials with target-congruent and target-incongruent orientation reports (268.56 *vs* 268.22 ms, *t*(8) = 0.29, *p* = 0.780).

Descriptively, saccade latencies and amplitudes showed small-scale variations across probe locations. To assess the significance of these variations, we performed two linear mixed-effects models in which we described the variance of amplitudes (*sacAmps*) or latencies (*sacLats*) with a fixed effect of probe location (*prbLoc*) and observer-specific, independent random effects for intercept and slope:

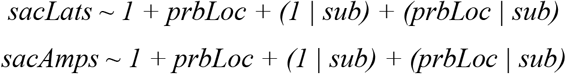

For this purpose, we recoded probe positions such that the foveal location was assigned the largest value. Values assigned to the remaining locations decreased linearly with eccentricity. The fixed effect of probe location was non-significant for saccade latencies (*t*(277) = −1.35, *p* = 0.177), with variations of less than 5 ms across probe locations. Probe location did affect saccade amplitudes, *t*(277) = −2.89, *p* = 0.004: amplitudes were shortest if the probe appeared in the fovea (*min* = 9.37 at *x* = 0 dva) and increased with probe eccentricity. These variations in saccade amplitude, though systematic, ranged within 0.17 dva (~5 pixels).

Subsequently, we investigated whether saccade amplitudes, saccade latencies, or their respective interaction with probe location had a meaningful impact on congruency effects (*HR_cong-incong_*). We added a random intercept of observer to account for inter-individual differences:

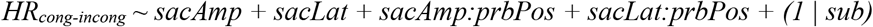

None of the main effects or interactions reached significance (all *t*s < 1.4, all *p*s > 0.16), suggesting that the demonstrated spatial profile of enhancement is independent of eye movement characteristics measured in our specific experimental design.

### Alternative function fits in Experiment 2

To allow for an asymmetric shape of spatial profiles within each observer, we fitted two-term Gaussian functions to individual-observer data:

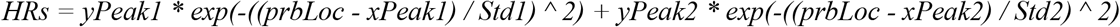

where *yPeak1, xPeak1* and *Std1* denote the y-coordinate of the peak, the x-coordinate of the peak and the standard deviation of the first Gaussian curve while *yPeak2, xPeak2* and *Std2* denote the equivalent parameters of the second Gaussian curve. The resulting profiles are plotted in **Supplementary Figure S1B**. This fitting approach yielded a mean absolute error of 0.7% for congruent and 1.0% for incongruent HRs and was therefore highly comparable to the more parsimonious Gaussian fit with vertical offset. A one-sample t-test on mean absolute errors revealed that the two fits were statistically indistinguishable, *t*(8) = −1.64, *p* = 0.139.

**Figure S1.**
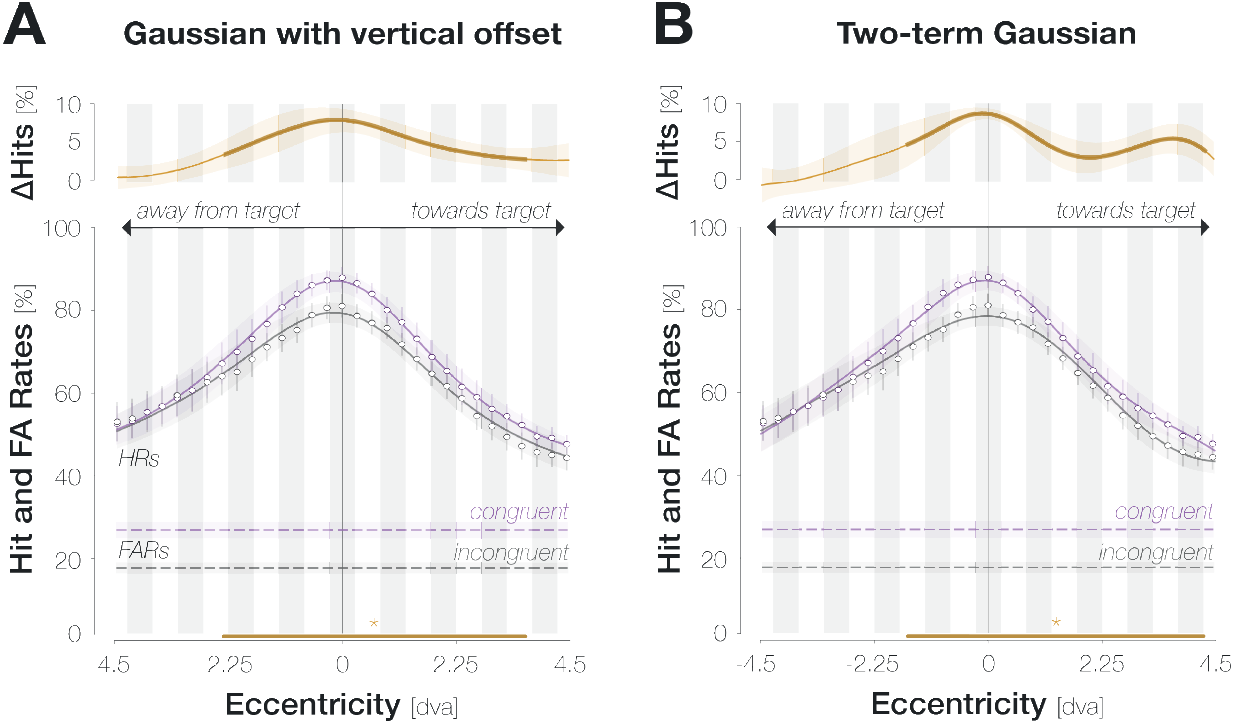
Gaussian functions with vertical offsets (**A**) and two-term Gaussian functions (**B**) yield comparable fits to spatial HR profiles. Conventions as in Figure 2D.

### Spatial development of noise content

Our design innately provides spatial resolution by allowing us to relate response behavior to background noise properties at any desired display location. We made use of this possibility in an exploratory analysis and evaluated if – despite observers’ explicit knowledge about the possible range of probe locations in Experiment 2 – FAs were primarily triggered by foveal orientation information. Specifically, we determined the energy around the reported orientation (target orientation ±22.5° for congruent FAs; non-target orientation ±22.5° for incongruent FAs) in the background noise along the axis of potential probe locations. Again, we collapsed all noise images that had appeared from the onset of the dynamic noise stream to saccade onset. Filter locations matched the 37 experimentally defined probe locations and were combined into 31 moving windows just like probe locations were. Before the moving window analysis, energy values were normalized to the mean and standard deviation of all filter responses for that specific SF*ori combination at that specific location. A movie displaying the spatial development of mean filter responses across spatial locations is provided as **Supplementary Movie 1**.

For both congruent and incongruent FAs, the energy of the reported orientation was highest in or close to the center of gaze (congruent: 0 dva; incongruent: 0.30 dva; **Figure S2**). Energy values for congruent FAs significantly exceeded zero in a symmetrical range from −1.80 to 1.80 dva (all *p*s < .041; bootstrapping, 10,000 repetitions). Energy values for incongruent FAs significantly exceeded zero at two locations: 0 and 0.30 dva, *p*s < .048. The difference between the congruent and incongruent profile, however, did not reach significance at any location, all *p*s > .068. Note that the relation between a possible sensitization to target-congruent orientations and mean filter responses is complex and likely asymmetrical for congruent and incongruent information: That is, congruent FAs may readily be triggered by noise images with weak target-like orientation content, whereas incongruent FAs may require strong target-incongruent signals. While this pattern of results would be a consequence of sensitization, it would counteract any difference in normalized energy for congruent compared to incongruent responses.

**Figure S2.**
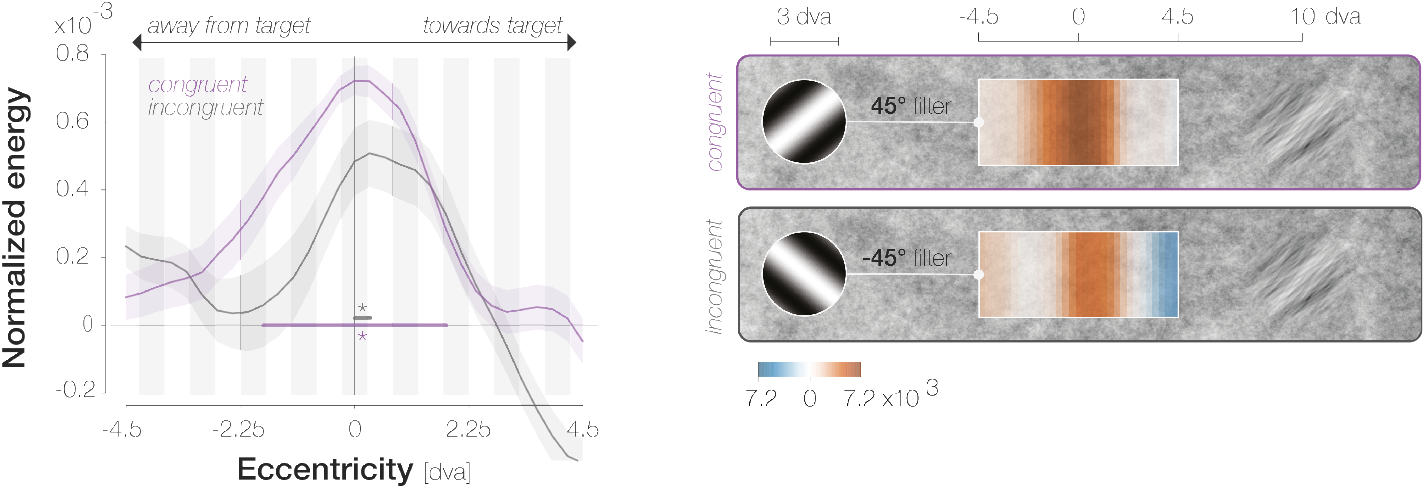
FAs are triggered by foveal orientation information. *Left*: Normalized energy around the reported orientation (+45° for congruent FAs, purple; −45° for incongruent FAs, gray) across spatial locations. Significance indicators highlight the spatial region in which the curve with corresponding color differs from zero. All further conventions follow those of Figure 2D. *Right*: Spatial profiles overlaid on the filtered region (drawn to scale with respect to the saccade target). Filters illustrate the relevant orientation in an example SF.

### Alignment of spatial profiles to the remapped location

Previous findings suggest that the location that visual information is predictively remapped to depends on the executed rather than intended vector of the impending saccade^[29]^. Due to both systematic^[38–40]^ and unsystematic variations in saccade endpoints, the predictively remapped location will more often than not differ from the pre-saccadic foveal location. To inspect if predictive sensitization is centered on the predictively remapped rather than the foveal location, we aligned probe positions to the remapped location of the target on an individual-trial level (**Figure S3**). Besides yielding flatter spatial profiles, this alignment reduced the spatial specificity of enhancement which now reached significance from −3.22 to 4.05 dva. These observations can be ascribed to the fact that the foveal location, in which both overall performance and enhancement is highest, contributes to different x-axis values in **Figure S3**. We conclude that an enhancement of saccade target features is invariably aligned to the pre-saccadic center of gaze – irrespective of the location attention is predictively remapped to on an individual-trial level.

**Figure S3.**
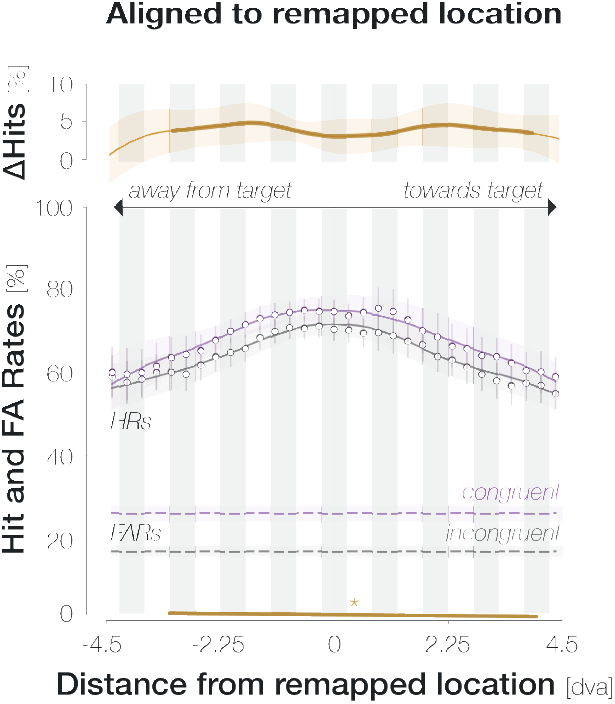
Foveal predictions are aligned to the center of gaze. Two-term Gaussian fit to the data plotted in S1 after probe locations were aligned to the predictively remapped target location rather than the pre-saccadic fixation. Conventions as in Figure 2D.

#### [Legend for Supplementary Movie 1]

*Movie S1*. FAs are primarily triggered by foveal orientation information. Filter responses of noise images underlying incongruent (**A**; the non-target orientation was reported) and congruent (**B**; the target orientation was reported) FAs at different spatial locations. The axis on top indicates the spatial range of filter locations (from −4.5 to 4.5 dva). The moving slider highlights the filter location corresponding to the currently displayed image. All further conventions as in Figure 2C. For both FA types, the energy around the reported orientation is particularly high in and around the foveal region.

